# Bovine H5N1 influenza virus binds poorly to human-type sialic acid receptors

**DOI:** 10.1101/2024.08.01.606177

**Authors:** Jefferson J.S. Santos, Shengyang Wang, Ryan McBride, Yan Zhao, James C. Paulson, Scott E. Hensley

**Author notes:** These authors contributed equally (Jefferson J.S. Santos, Shengyang Wang, Ryan McBride).

## Abstract

Clade 2.3.4.4b highly pathogenic H5N1 avian influenza (HPAI) viruses started circulating widely in lactating dairy cattle in the United States at the end of 2023. Avian influenza viruses enter cells after binding to glycan receptors with terminally linked α2-3 sialic acid, whereas human influenza viruses typically bind to glycan receptors terminally linked α2-6 sialic acid in the upper respiratory tract. Here, we evaluated the receptor binding properties of hemagglutinin (HA) trimers from a clade 2.3.4.4b avian isolate (A/American Wigeon/South Carolina/22-000345-001/2021) and a cattle isolate (A/dairy cattle/Texas/24-008749-002-v/2024). Using two different methods, we found that both of the 2.3.4.4b H5s bound efficiently to glycan receptors with terminally linked α2-3 sialic acid with no detectable binding to glycan receptors with terminally linked α2-6 sialic acid. Our data suggest that clade 2.3.4.4b H5N1 viruses bind poorly to human receptors. It will be important to continue evaluating receptor binding properties of these viruses as they evolve in cattle.

---

Highly pathogenic H5N1 avian influenza (HPAI) viruses started circulating in lactating dairy cattle in the United States at the end of 2023^1^ and these viruses are now rapidly spreading between cows^2^. Eisfeld et al. reported that a clade 2.3.4.4b H5N1 virus from this cattle outbreak binds to α2-6 linked sialylglycopolymers coated to microtiter plates^3^. This is an important finding since human influenza viruses bind efficiently to glycan receptors with terminally linked α2-6 sialic acid that are abundant in the upper respiratory tract of humans, whereas avian influenza viruses typically bind to glycan receptors with terminally linked α2-3 sialic acid. Moreover, acquisition of human-type receptor specificity is believed to be required for efficient transmission of influenza virus in humans and is considered a risk factor for the emergence of a new pandemic virus^4^.

In cattle, H5N1 replicates efficiently in the mammary gland^2^, which expresses both α2-3 and α2-6 linked sialic acids^5^. This together with the results of Eisfeld et al.^3^ suggest that the 2.3.4.4b H5 viruses may be evolving to more efficiently bind to “human-type” α2-6 linked sialic acids. To directly compare the receptor specificity of the H5N1 virus circulating in birds with the bovine H5N1 virus we created soluble recombinant hemagglutinin (HA) trimers from the A/American Wigeon/South Carolina/22-000345-001/2021 (A/South Carolina/2021) and A/dairy cattle/Texas/24-008749-002-v/2024 (A/Cattle/Texas/2024) H5N1 viruses. A/South Carolina/2021 is a World Health Organization candidate vaccine virus isolated prior to the cattle outbreak and the A/Cattle/Texas/2024 virus was isolated from a dairy cow in 2024. The A/Cattle/Texas/2024 HA has only 2 amino acid substitutions (L122Q and T199I; H3 numbering) relative to the A/South Carolina/2021 HA. We assessed the receptor binding properties of the A/South Carolina/2021 and A/Cattle/Texas/2024 HAs using two complementary assays.

First, we tested HA binding to biotinylated glycans adsorbed to high-binding capacity streptavidin-coated plates. Glycans contained terminal sequences with either the avian-type (NeuAcα2-3Galβ1-4GlcNAc) or human-type (NeuAcα2-6Galβ1-4GlcNAc) receptor, including three linear glycans with one, two or three N-acetyl-lactosamine (LN, Galβ1-4GlcNAc) repeats (3SLN1/2/3-L or 6SLN1/2/3-L), and three biantennary N-linked glycans displaying the same sequences on both branches (3SLN1/2/3-N or 6SLN1/2/3-N); **Fig. 1a**). Glycans were displayed at a consistent density by loading the plate with an excess of biotinylated glycans to saturate the streptavidin-coated surface and we tested different dilutions of HA. Glycan loading was verified using serially diluted Maackia amurensis agglutinin (MAA) and Sambucus nigra agglutinin (SNA), which bind respectively to α2-3 and α2-6 linked sialic acids (**Fig. 1b**). The A/South Carolina/2021 H5 HA exhibited strict avian-type receptor specificity, binding all linear and N-linked glycans terminated in α2-3 linked sialic acids, with no detectable binding to the glycans containing α2-6 linked sialic acids. The A/Cattle/Texas/2024 HA similarly exhibited strict avian-type specificity. Both the A/South Carolina/2021 and A/Cattle/Texas/2024 H5s bound to glycans with α2-3 linked sialic acids with >100 fold higher avidity than to the glycans with α2-6 sialic acids (**Fig. 1c**).

**Fig. 1.**
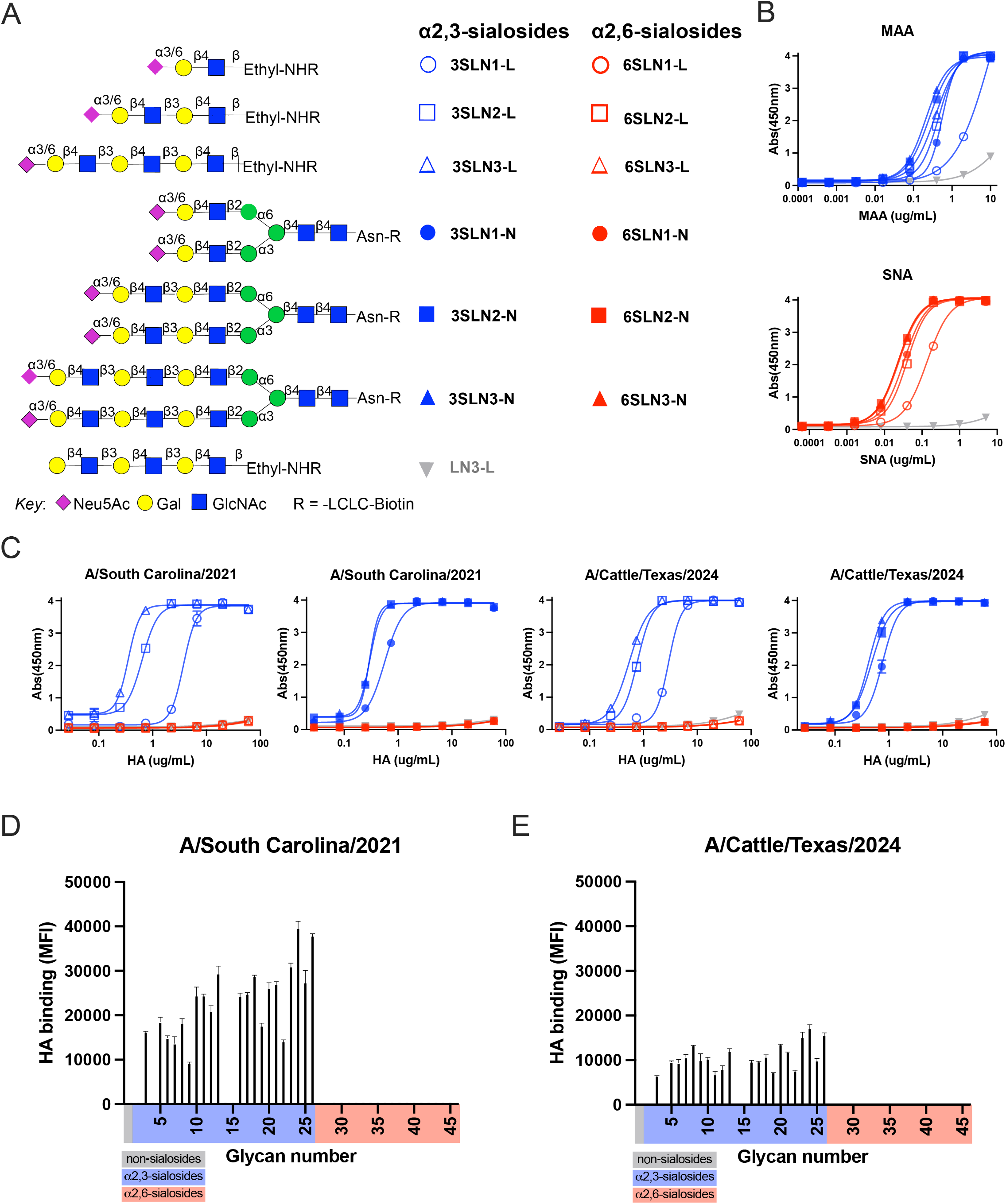
Bovine H5N1 influenza HA binds retains high specificity for avian-type sialic acid receptors. Glycan binding analysis of avian and cattle clade 2.3.4.4b H5 HAs. Panel A shows list of the biotinylated glycans analyzed in an ELISA based format, with terminal α2-3 or α2-6 linked sialic acids representing avian-type or human-type receptors, respectively. Panel B shows binding of Maackia amurensis agglutinin (MAA) and Sambucus nigra agglutinin (SNA) to assess loading for glycans with α2-3 and α2-6 linked sialic acids, respectively. Panel C displays results for A/South Carolina/2021 and A/Cattle/Texas/2024 H5 HA binding to terminally sialylated linear (L) and biantennary N-linked (N) glycans. Panel D Glycan microarray analysis of (A/South Carolina/2021) and E (A/Cattle/Texas/2024) H5 HAs. The array contains a library of glycans comprising control glycans with no sialic acid (grey), α2-3 sialosides (blue) and α2-6 sialosides (red). Shown is mean and standard error of the mean SEM of 4 replicates. A complete list of glycan structures elaborated on the array are found in Extended Table 1.

Next, we tested HA binding to a sialoside glycan microarray containing avian-type and human-type receptors representing the structural diversity of natural glycans terminated with α2-3 or α2-6 linked sialic acids^6^. As shown in **Fig. 1d,e** both the A/South Carolina/2021 and A/Cattle/Texas/2024 HAs bound a wide variety of the sialic acid α2-3 terminated glycans, with no significant binding to sialic acid α2-6 terminated glycans. Thus, the two assays show no significant change in the specificity of the A/South Carolina/2021 and A/Cattle/Texas/2024 HAs for avian-type receptors or the breadth of structural diversity represented in the glycan microarray.

Our results suggest that the bovine H5N1 virus has maintained strict specificity for binding to avian-type receptors, as seen by others using glycan microarray to assess receptor specificity^7^. Results of Eisfeld et al.^3^ suggest that human-type receptor specificity is evident for both an avian H5N1 virus and a dairy cattle H5N1 virus (A/dairy cattle/New Mexico/A240920343-93/2024). Differences might be attributed to the fact that whole virus with higher HA valency was used, and/or differences in the assay using sialoside polymers with avian- and human-type receptors. Previous studies suggest that several HA mutations are needed to substantially change the specificity of the H5N1 virus to human-type receptor specificity^4^. Continued surveillance of H5N1 in cattle and other species is warranted to track the potential of the virus to acquire human-type receptor specificity which would increase the risk of its emergence of a pandemic virus.

## Methods

### Recombinant HA proteins

Soluble recombinant HA trimers were produced as previously described^8^. HA mammalian expression plasmids were created with codon-optimized full-length HA sequences. The HA transmembrane domain was replaced with a FoldOn trimerization domain from T4 fibritin, the AviTag for site-specific biotinylation, and a hexahistidine tag for affinity purification. Plasmids were transfected into 293F suspension cells and supernatant from transfected was harvested and clarified by centrifugation 4 days later. HA proteins were purified using Ni-NTA agarose resin (Qiagen) and gravity flow chromatography columns (Bio-Rad). HA was buffer-exchanged and concentrated using centrifugal filter units with a 30 kDa molecular weight cutoff (Millipore).

### Glycan ELISA

HAs were assessed for their avidity for binding synthetic biotinylated glycans as previously described^9,10^. Briefly, streptavidin-coated high binding capacity 384-well plates (Pierce) were incubated with 1.8 μM biotinylated glycan in PBS at 4 °C overnight. Plates were rinsed with PBS containing 0.05% Tween-20 (PBS-T) 5 times to remove excess glycans, blocked with 1% bovine serum albumin (BSA) in PBS (BSA/PBS) buffer containing 0.6 μM desthiobiotin at room temperature for 2 h. Plates were then washed with PBS-T 5 times and used without further processing. Purified His-tagged-HAs were premixed with anti-His mouse IgG2a antibody (Biolegend, cat no. 362616) and HRP-conjugated goat anti-mouse IgG (H + L) secondary antibody (Invitrogen, cat no. G21040) in a 4:2:1 ratio (w/w/w) in BSA/PBS buffer and incubated on ice for 30 min. The HA complex was then subjected to 3-fold serial dilutions, added to wells of glycan-coated plates and incubated at room temperature for 2 h. Plates then washed and developed by adding 3,3′,5,5′ tetramethylbenzidine (TMB, Sigma-Aldrich, cat no. T0440) peroxidase substrate to wells for 15 min at room temperature. Reaction was quenched with 2 M sulfuric acid and absorbance at 450 nm was detected using a BioTek Synergy H1 microplate reader (Agilent). The assay was performed in duplicate. *Maackia amurensis* agglutinin (MAA) and the *Sambucus nigra* agglutinin (SNA) were used as positive control for SAα2,3Gal and SAα2,6Gal binding, respectively.

### Glycan array analyses

Glycans microarrays were produced, and analysis was performed as described previously^6,9^. Purified His-tagged-HAs were premixed with anti-His mouse IgG2a antibody (Invitrogen, MA1-21315) and HRP-conjugated goat anti-mouse IgG (H + L) secondary antibody(Invitrogen, A001) in PBS-T in a 4:2:1 ratio and incubated on ice for 15 min. Pre-complexed HAs were incubated on the microarray surface for 60 min in a humidified chamber at room temperature. Slides were washed twice in PBS-T, PBS, then water, and dried until arrays were scanned using an Innoscan 1100AL microarray scanner (Innopsys). Slides contained six replicates of each glycan. Replicates with the highest and lowest fluorescence intensity were excluded, and the fluorescence intensity of the remaining four replicates was used to obtain the mean and standard error of the mean (SEM).

## Data Availability

No code was used in this study. Source data are available by request from the corresponding authors.

## Author Contribution

J.J.S.S., S.W., R.M. Y.Z. completed experiments and analyzed data. J.C.P. and S.E.H. supervised experiments. J.J.S.S. wrote the 1^st^ draft of the manuscript and all authors edited the manuscript.

## Acknowledgments

This project has been funded in part with Federal funds from the National Institute of Allergy and Infectious Diseases, National Institutes of Health (NIAID), Department of Health and Human Services, under Contract No. 75N93021C00015 and NIAID grant number AI114730.

## Conflict of Interest

S.E.H reports receiving consulting fees from Sanofi, Pfizer, Lumen, Novavax, and Merck. The authors declare no other competing interests.

