## Extended Table 1 for "Bovine H5N1 influenza virus binds poorly to human-type sialic acid receptors"

Extended Table 1. Glycans on the glycan microarray.

| ChartNo | Catalog Number | Structure | Symbol Structure |
|---|---|---|---|
| 1 | M040 | Galβ(1-4)-GlcNAcβ-ethyl-NH2 | 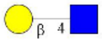 |
| 2 | M009 | Galβ(1-4)-GlcNAcβ(1-2)-Manα(1-3)-[Galβ(1-4)-GlcNAcβ(1-2)-Manα(1-6)]-Manβ(1-4)-GlcNAcβ(1-4)-GlcNAcβ-Asn-$NH_2$ | 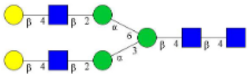 |
| 3 | SW32 | NeuAcα(2-3)-Galβ(1-4)-GlcNAcβ-ethyl-$NH_2$ | 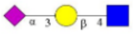 |
| 4 | SW33 | NeuAcα(2-3)-Galβ(1-4)-GlcNAcβ(1-3)-Galβ(1-4)-GlcNAcβ-ethyl-$NH_2$ | 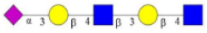 |
| 5 | SW34 | NeuAcα(2-3)-Galβ(1-4)-GlcNAcβ(1-3)-Galβ(1-4)-GlcNAcβ(1-3)-Galβ(1-4)-GlcNAcβ-ethyl-NH2 | 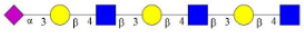 |
| 6 | M045 | NeuAcα(2-3)-Galβ(1-3)-GalNAca-Thr-$NH_2$ | 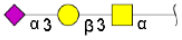 |
| 7 | M120 | NeuAcα(2-3)-Galβ(1-4)-GlcNAcβ(1-3)-Galβ(1-3)-GalNAca-Thr-NH2 | 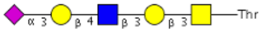 |
| 8 | M128 | NeuAcα(2-3)-Galβ(1-4)-GlcNAcβ(1-3)-Galβ(1-4)-GlcNAcβ(1-3)-Galβ(1-3)-GalNAca-Thr-NH2 | 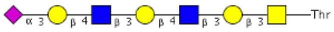 |
| 9 | M142 | NeuAcα(2-3)-Galβ(1-4)-GlcNAcβ(1-3)-Galβ(1-4)-GlcNAcβ(1-3)-Galβ(1-4)-GlcNAcβ(1-3)-Galβ(1-3)-GalNAca-Thr-NH2 | 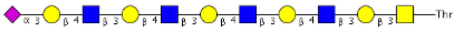 |
| 10 | M143 | NeuAcα(2-3)-Galβ(1-4)-GlcNAcβ(1-3)--Galβ(1-4)-GlcNAcβ(1-3)-Galβ(1-4)-GlcNAcβ(1-3)-Galβ(1-4)-GlcNAcβ(1-3)-Galβ(1-3)-GalNAca-Thr-NH2 | 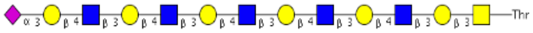 |
| 11 | M050 | NeuAcα(2-3)-Galβ(1-4)-GlcNAcβ(1-6)-[Galβ(1-3)]-GalNAca-Thr-$NH_2$ | 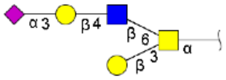 |
| 12 | M053 | NeuAcα(2-3)-Galβ(1-4)-GlcNAcβ(1-3)-Galβ(1-4)-GlcNAcβ(1-6)-[Galβ(1-3)]-GalNAca-Thr-$NH_2$ | 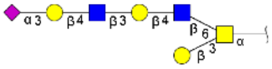 |

| | | | |
|---|---|---|---|
| **13** | M202 | NeuAcα(2-3)-Galβ(1-4)-GlcNAcβ(1-3)-Galβ(1-4)-GlcNAcβ(1-3)-Galβ(1-4)-GlcNAcβ(1-6)-[Galβ(1-3)]-GalNAca-Thr-$NH_2$ | 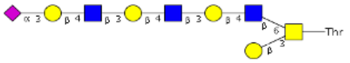  |
| **14** | M152 | NeuAcα(2-3)-Galβ(1-4)-GlcNAcβ(1-3)-Galβ(1-4)-GlcNAcβ(1-3)-Galβ(1-4)-GlcNAcβ(1-3)-Galβ(1-4)-GlcNAcβ(1-6)-[Galβ(1-3)]-GalNAca-Thr-$NH_2$ | 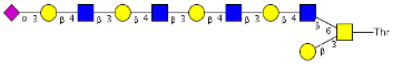  |
| **15** | M149 | NeuAcα(2-3)-Galβ(1-4)-GlcNAcβ(1-3)-Galβ(1-4)-GlcNAcβ(1-3)-Galβ(1-4)-GlcNAcβ(1-3)-Galβ(1-4)-GlcNAcβ(1-3)-Galβ(1-4)-GlcNAcβ(1-6)-[Galβ(1-3)]-GalNAca-Thr-$NH_2$ | 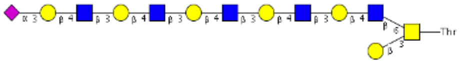  |
| **16** | SW07 | NeuAcα2-3Galβ1-4GlcNAcβ1-2Manα1-3(NeuAcα2-3Galβ1-4GlcNAcb1-2Manα1-6)Manβ1-4GlcNAcβ1-4GlcNAc-AsnGly | 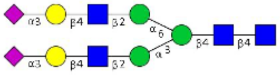 |
| **17** | SW08 | NeuAca2-3Galb1-4GlcNAcb1-3Galb1-4GlcNAcb1-2Mana1-3(NeuAca2-3Galb1-4GlcNAcb1-3Galb1-4GlcNAcb1-2Mana1-6)Manb1-4GlcNAcb1-4GlcNAc-AsnGly | 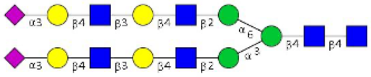 |
| **18** | SW09 | (NeuAca2-3Galb1-4GlcNAcb1-3Galb1-4GlcNAcb1-3Galb1-4GlcNAcb1-2Mana1-6)NeuAca2-3Galb1-4GlcNAcb1-3Galb1-4GlcNAcb1-3Galb1-4GlcNAcb1-2Mana1-3Manb1-4GlcNAcb1-4GlcNAc-AsnGly | 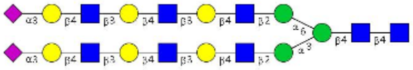 |
| **19** | SW10 | NeuAca2-3Galb1-4GlcNAcb1-2Mana1-3(NeuAca2-3Galb1-4GlcNAcb1-2Mana1-6)Manb1-4GlcNAcb1-4(Fuca1-6)GlcNAc-AsnGly | 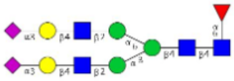 |
| **20** | SW11 | NeuAca2-3Galb1-4GlcNAcb1-3Galb1-4GlcNAcb1-2Mana1-3(NeuAca2-3Galb1-4GlcNAcb1-3Galb1-4GlcNAcb1-2Mana1-6)Manb1-4GlcNAcb1-4(Fuca1-6)GlcNAc-AsnGly | 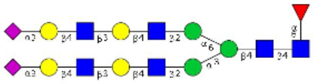 |
| **21** | SW12 | NeuAca2-3Galb1-4GlcNAcb1-3Galb1-4GlcNAcb1-3Galb1-4GlcNAcb1-2Mana1-3(NeuAca2-3Galb1-4GlcNAcb1-3Galb1-4GlcNAcb1-3Galb1-4GlcNAcb1-2Mana1-6)Manb1-4GlcNAcb1-4GlcNAc-AsnGly | 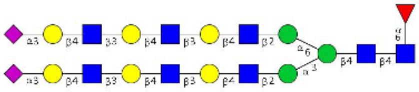 |
| **22** | M002 | NeuAca(2-3)-Galβ(1-4)-[Fuca(1-3)]-GlcNAcβ-propyl-$NH_2$ | 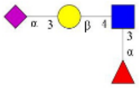 |
| **23** | M022 | NeuAca(2-3)-Galβ(1-4)-[Fuca(1-3)]-GlcNAcβ(1-3)-Galβ(1-4)-[Fuca(1-3)]-GlcNAcβ-ethyl-$NH_2$ | 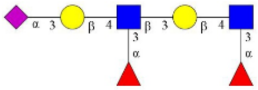 |

| | | | |
|---|---|---|---|
| 24 | M015 | NeuAca(2-3)-Galβ(1-4)-[Fuca(1-3)]-GlcNAcβ(1-3)-Galβ(1-4)-[Fuca(1-3)]-GlcNAcβ(1-3)-Galβ(1-4)-[Fuca(1-3)]-GlcNAcβ-ethyl-$NH_2$ | 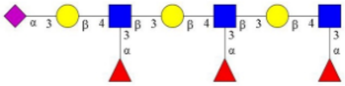 |
| 25 | M206 | NeuAca(2-3)-Galβ(1-4)-[Fuca(1-3)]-GlcNAcβ(1-3)-Galβ(1-4)-[Fuca(1-3)]-GlcNAcβ(1-3)-Galβ(1-4)-[Fuca(1-3)]-GlcNAcβ(1-3)-Galβ(1-3)-GalNAca-Thr-NH2 | 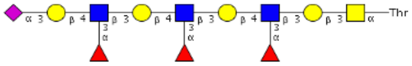  |
| 26 | M147 | NeuAca(2-3)-Galβ(1-4)-[Fuca(1-3)]-GlcNAcβ(1-3)-Galβ(1-4)-[Fuca(1-3)]-GlcNAcβ(1-3)-Galβ(1-4)-[Fuca(1-3)]-GlcNAcβ(1-3)-GalNAca-Thr-NH2 | 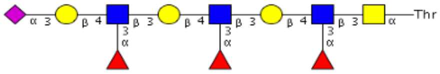  |
| 27 | M215 | NeuAcα(2-6)-Galβ(1-4)-(6S)GlcNAcβ-ethyl-NH2 | 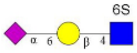  |
| 28 | M003 | NeuAca(2-6)-Galβ(1-4)-6-O-sulfo-GlcNAcβ-propyl-$NH_2$ | 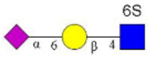  |
| 29 | SW29 | NeuAca(2-6)-Galβ(1-4)-GlcNAcβ-ethyl-$NH_2$ | 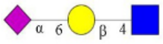 |
| 30 | SW30 | NeuAca(2-6)-Galβ(1-4)-GlcNAcβ(1-3)-Galβ(1-4)-GlcNAcβ-ethyl-$NH_2$ | 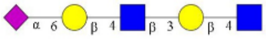 |
| 31 | SW31 | NeuAca(2-6)-Galβ(1-4)-GlcNAcβ(1-3)-Galβ(1-4)-GlcNAcβ(1-3)-Galβ(1-4)-GlcNAcβ-ethyl-$NH_2$ | 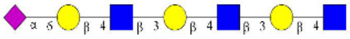 |
| 32 | M121 | NeuAcα(2-6)-Galβ(1-4)-GlcNAcβ(1-3)-Galβ(1-3)-GalNAca-Thr-NH2 | 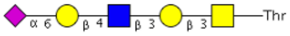  |
| 33 | M129 | NeuAcα(2-6)-Galβ(1-4)-GlcNAcβ(1-3)-Galβ(1-4)-GlcNAcβ(1-3)-Galβ(1-3)-GalNAca-Thr-NH2 | 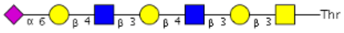  |
| 34 | M154 | NeuAcα(2-6)-Galβ(1-4)-GlcNAcβ(1-3)-Galβ(1-4)-GlcNAcβ(1-3)-Galβ(1-4)-GlcNAcβ(1-3)-Galβ(1-3)-GalNAca-Thr-NH2 | 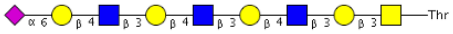  |

| | | | |
|---|---|---|---|
| **35** | M135 | NeuAcα(2-6)-Galβ(1-4)-GlcNAcβ(1-3)-Galβ(1-4)-GlcNAcβ(1-3)-Galβ(1-4)-GlcNAcβ(1-3)-Galβ(1-4)-GlcNAcβ(1-3)-Galβ(1-3)-GalNAca-Thr-NH2 | 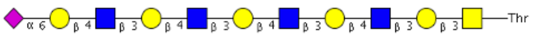 |
| **36** | M051 | NeuAcα(2-6)-Galβ(1-4)-GlcNAcβ(1-6)-[Galβ(1-3)]-GalNAca-Thr-$NH_2$ | 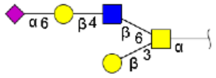 |
| **37** | M054 | NeuAcα(2-6)-Galβ(1-4)-GlcNAcβ(1-3)-Galβ(1-4)-GlcNAcβ(1-6)-[Galβ(1-3)]-GalNAca-Thr-$NH_2$ | 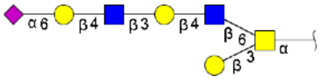 |
| **38** | M201 | NeuAcα(2-6)-Galβ(1-4)-GlcNAcβ(1-3)-Galβ(1-4)-GlcNAcβ(1-3)-Galβ(1-4)-GlcNAcβ(1-6)-[Galβ(1-3)]-GalNAca-Thr-$NH_2$ | 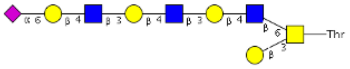 |
| **39** | M159 | NeuAcα(2-6)-Galβ(1-4)-GlcNAcβ(1-3)-Galβ(1-4)-GlcNAcβ(1-3)-Galβ(1-4)-GlcNAcβ(1-3)-Galβ(1-4)-GlcNAcβ(1-6)-[Galβ(1-3)]-GalNAca-Thr-$NH_2$ | 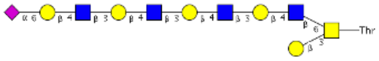 |
| **40** | M157 | NeuAcα(2-6)-Galβ(1-4)-GlcNAcβ(1-3)-Galβ(1-4)-GlcNAcβ(1-3)-Galβ(1-4)-GlcNAcβ(1-3)-Galβ(1-4)-GlcNAcβ(1-3)-Galβ(1-4)-GlcNAcβ(1-6)-[Galβ(1-3)]-GalNAca-Thr-$NH_2$ | 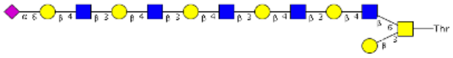 |
| **41** | SW01 | NeuAca2-6Galb1-4GlcNAcb1-2Mana1-3(NeuAca2-6Galb1-4GlcNAcb1-2Mana1-6)Manb1-4GlcNAcb1-4GlcNAc-AsnGly | 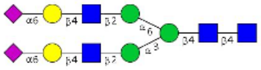 |
| **42** | SW02 | NeuAca2-6Galb1-4GlcNAcb1-3Galb1-4GlcNAcb1-2Mana1-3(NeuAca2-6Galb1-4GlcNAcb1-3Galb1-4GlcNAcb1-2Mana1-6)Manb1-4GlcNAcb1-4GlcNAc-AsnGly | 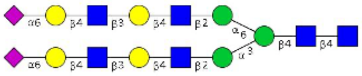 |
| **43** | SW03 | NeuAca2-6Galb1-4GlcNAcb1-3Galb1-4GlcNAcb1-3Galb1-4GlcNAcb1-2Mana1-3(NeuAca2-6Galb1-4GlcNAcb1-3Galb1-4GlcNAcb1-3Galb1-4GlcNAcb1-2Mana1-6)Manb1-4GlcNAcb1-4GlcNAc-AsnGly | 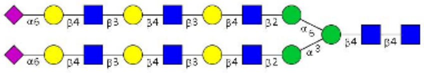 |
| **44** | SW04 | NeuAca2-6Galb1-4GlcNAcb1-2Mana1-3(NeuAca2-6Galb1-4GlcNAcb1-2Mana1-6)Manb1-4GlcNAcb1-4(Fuca1-6)GlcNAc-AsnGly | 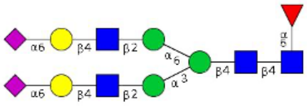 |

| | | | |
|---|---|---|---|
| **45** | SW05 | NeuAca2-6Galb1-4GlcNAcb1-3Galb1-4GlcNAcb1-2Mana1-3(NeuAca2-6Galb1-4GlcNAcb1-3Galb1-4GlcNAcb1-2Mana1-6)Manb1-4GlcNAcb1-4(Fuca1-6)GlcNAc-AsnGly | 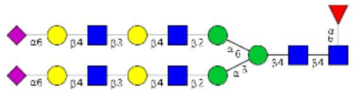 |
| **46** | SW06 | NeuAca2-6Galb1-4GlcNAcb1-3Galb1-4GlcNAcb1-3Galb1-4GlcNAcb1-2Mana1-3(NeuAca2-6Galb1-4GlcNAcb1-3Galb1-4GlcNAcb1-3Galb1-4GlcNAcb1-2Mana1-6)Manb1-4GlcNAcb1-4GlcNAc-AsnGly | 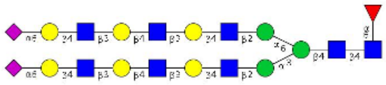 |
